# Homologous Recombination Shapes the Architecture and Evolution of Bacterial Genomes

**DOI:** 10.1101/2024.05.31.596828

**Authors:** Ellis L. Torrance, Awa Diop, Louis-Marie Bobay

**Affiliations:** Dept. of Biology, University of North Carolina Greensboro, Greensboro, NC 27412; Systems Biology Dept., Sandia National Laboratories, Livermore, CA 94551; Dept. of Biological Sciences, North Carolina State University, Raleigh, NC 27695

**Author notes:** E.L.T., A.D., and L-M. B. designed, performed research, and analyzed data in this publication. E.L.T. and L-M.B. wrote this paper and A.D. reviewed and edited it.

## Abstract

Homologous recombination is a key evolutionary force that varies considerably across bacterial species. However, how the landscape of homologous recombination varies across genes and within individual genomes has only been studied in a few species. Here, we used Approximate Bayesian Computation to estimate the recombination rate along the genomes of 145 bacterial species. Our results show that homologous recombination varies greatly along bacterial genomes and shapes many aspects of genome architecture and evolution. The genomic landscape of recombination presents several key signatures: rates are highest near the origin of replication in most species, patterns of recombination generally appear symmetrical in both replichores (*i.e.* replicational halves of circular chromosomes) and most species have genomic hotpots of recombination. Furthermore, many closely related species share conserved landscapes of recombination across orthologs indicating that recombination landscapes are conserved over significant evolutionary distances. We show evidence that recombination drives the evolution of GC-content through increasing the effectiveness of selection and not through biased gene conversion, thereby contributing to an ongoing debate. Finally, we demonstrate that the rate of recombination varies across gene function and that many hotspots of recombination are associated with adaptive and mobile regions often encoding genes involved in pathogenicity.

## Introduction

Homologous recombination is a major force shaping genome evolution across bacteria. This mechanism promotes the exchange of alleles between homologous sequences akin to the process of gene conversion in Eukaryotes. The rates of recombination vary extensively across species, but several studies have shown that recombination rates vary across genomic regions as well^1–3^. In addition to the overall fluctuations of recombination rates, some genomic regions present very low rates of recombination (*i.e*., coldspots) while others show particularly high rates of recombination (*i.e*., hotspots)^4^. The mechanisms shaping these variations in recombination rate along the genome are not known. Some regions might be more recombinogenic by presenting an easier access for incoming DNA or by containing sequence motifs that stimulate recombination (*e.g*., Chi motifs)^5,6^. Alternatively, selection may be shaping the patterns of recombination across genomic regions. It has been hypothesized that hotspots of recombination play an important role in the ability of bacteria to adapt to environmental selective pressures^7^. However, the patterns of homologous recombination have only been characterized in the genomes of a few species^4,5,7,8^.

Thus, it remains largely unknown how variations in homologous recombination contribute to driving the evolution and adaptation of bacterial genomes. In addition, to what extent these genomic landscapes of recombination are conserved after species diverge from one another remains to be determined.

In Eukaryotes, the genomic landscapes of recombination have been extensively studied and hotspots of recombination have been found to be associated with adaptive phenotypes^9^. In bacteria however, varying recombination rates across genomes have not been thoroughly described^3,4,7,8^. Previous studies have shown that signatures of recombination are lower in core genes with housekeeping functions and higher in genes associated with virulence and defense functions^4,7,8,10^. Interestingly, hotspots of recombination have been observed to be flanking mobile elements such as SCC (Staphylococcal cassette chromosome) which is associated with methicillin resistance in *Staphylococcus aureus* (MRSA)^7,8,11^. Hotspots of recombination have also been found near the origin of replication (Ori) in *S. aureus*, but this trend has not been observed in other species^7,8^. Moreover, genes flanking mobile elements and clusters of accessory genes have been estimated to recombine twice as frequently as non-flanking genes across bacterial species^12^.

Bacterial chromosomes present various levels of organization, and those can impose mechanistic and selective constraints on recombination. Replication proceeds bidirectionally starting at the Ori^13^. In circular chromosomes, the two replichores present similar lengths and progress synchronously from Ori to the terminus of replication (Ter). A mutation accumulation study has shown that mutation frequency follows a wave-like pattern, symmetrical in the two replichores, which has been hypothesized to be the result of replication timing and its interruption^14^. Because DNA synthesis proceeds necessarily from 5’ to 3’, one strand is synthetized continuously in the same direction as the replication fork (*i.e*., the leading strand), while the other strand is synthetized discontinuously (*i.e*., the lagging strand). Thus, the initiation of DNA synthesis is delayed for the lagging strand and its template strand remains single-stranded for longer periods of time. Due to the higher mutagenic nature of single-stranded DNA, it has been suggested that this asymmetry in DNA replication causes higher mutation rates on the lagging strand relative to the leading strand^15^. However, it has been shown that this asymmetry could in fact be driven by the stronger selective constraints acting on the leading strand, due to the higher prevalence of essential and highly expressed genes on this strand^13^. Finally, accessory genes are typically organized into gene clusters such as pathogenicity islands, which are often horizontally exchanged between strains^13,16^. The insertion of these transferred clusters of accessory genes can be mediated by non-homologous recombination, site-specific recombination, or by homologous recombination at flanking core genes^17–19^.

Here, we used a new approach based on Approximate Bayesian Computation (ABC) to estimate the rates of homologous recombination along bacterial genomes. We adapted our tool *recABC*^20^ to infer the recombination rate of thousands of individual core genes across the genomes of 145 bacterial species and one archaeon. We compared the rates of recombination in the context of their chromosomal location and functions to determine whether homologous recombination rate varies with any appreciable patterns across species. Using a robust dataset of >200,000 core genes, we observed that homologous recombination varies extensively across the genome of most bacteria. We further found evidence that these variations are linked to gene function and chromosome structure, indicating that selection is shaping patterns of recombination. We detected the presence of many hotspots and coldspots of recombination across species. We did not identify any strand-specific biases in homologous recombination or any strong indication that recombination rate is elevated in genes directly flanking accessory regions. However, we did observe a significant relationship between homologous recombination and GC-content as well as signatures of selection.

## Results

### Homologous Recombination Rate Varies across Bacterial Core Genes

In a recent study, we developed and applied a novel approach to estimate recombination rate across bacteria using ABC^20^. Briefly, our tool generates forward-in-time simulations of genome populations evolved under various rates of recombination (*n*=500,000 simulations per species). Our approach selects the simulations that best recapitulate genomic signatures of recombination observed in the real genomes through comparison of summary statistics with ABC. We utilized the summary statistics of individual core gene alignments to estimate recombination rate for each gene using the simulated population of sequences (Fig. 1). For each core gene, we measured the effective rate of recombination *r/m*, which represents the number of times alleles have been exchanged by recombination (*r*) relative to the number of alleles introduced by mutation (*m*).

**Figure 1.**
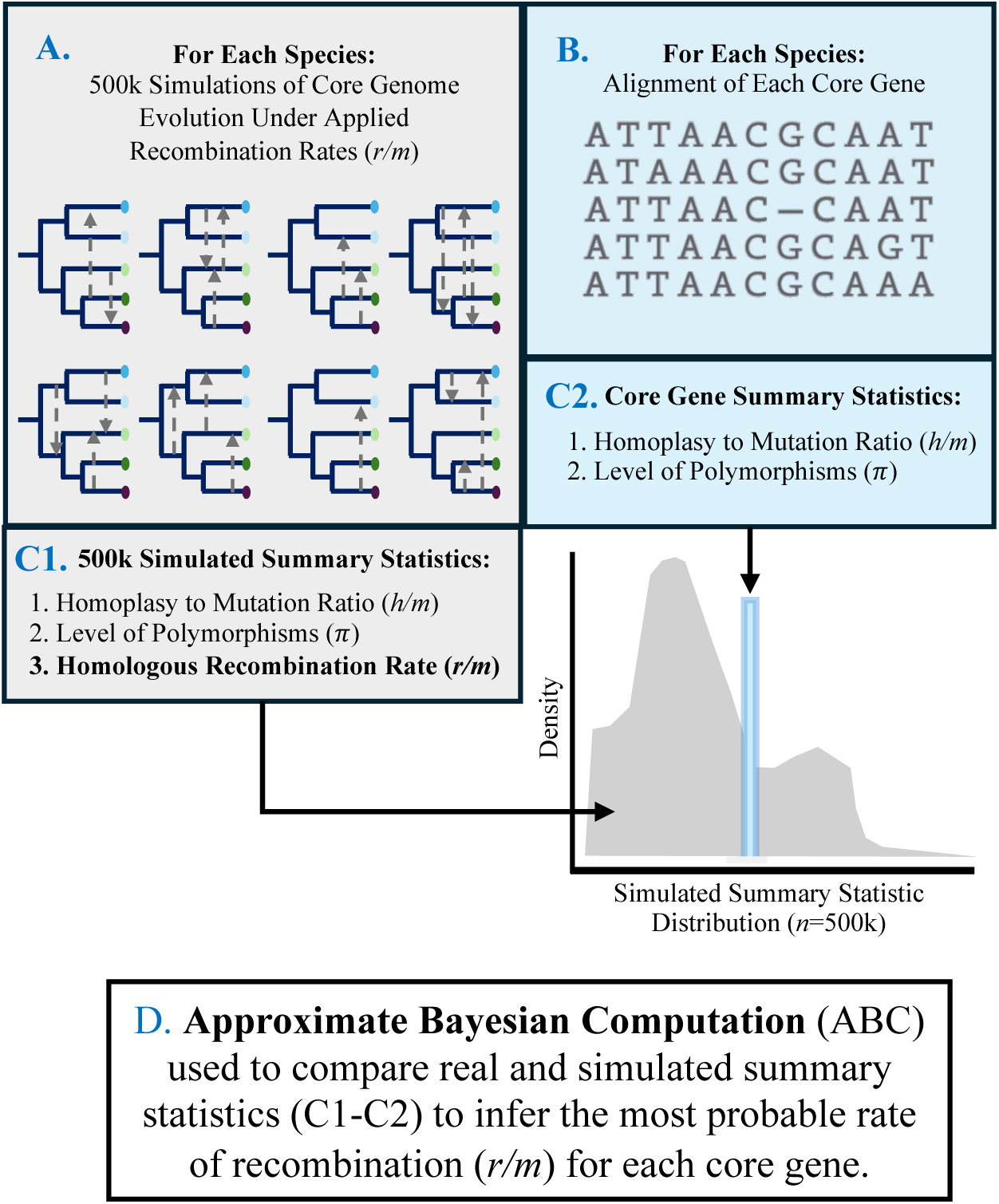
Description of the method used to infer the rates of homologous recombination (*recABC*) across the core genes of 145 bacteria and one archaeal species adapted from Torrance *et al.* (2024)^20^. A) For each species, core genes are aligned and concatenated and a maximum likelihood phylogenetic tree is built from the alignment of the core genome. Using the nucleotide length and GC-content of the core genome alignment from the real species, a single ancestral genome was randomly generated to initiate each simulation. This ancestral genome was then evolved in a forward-in-time simulation following the phylogenetic topology of the real species and its transition transversion bias under varied recombination rates (*Rho*) and recombination tract lengths (*delta*) using *CoreSimul* ^21^. The corresponding effective recombination rate *r/m* was also computed from these parameters during the simulations. A total of 500,000 simulations were generated for each species. B) An alignment was generated for each core gene within each species. C1) Two summary statistics are computed: *i*) the ratio of homoplasy to mutation (*h/m*), and *ii*) the average nucleotide diversity (*Pi*) computed from the core genome alignment of the real species. C2) The same summary statistics (*h/m, Pi*) are calculated for each of the 500k simulations. D) Approximate Bayesian Computation (ABC) is then used to compare the summary statistics from the real species to the distribution of the same statistics generated by simulation under known recombination rates (the prior distribution in grey). The simulations whose statistics most closely match the summary statistics from the real species are selected using ABC (the posterior distribution in blue) with a tolerance threshold of 0.01% (*n*=50). The average rate of recombination of the posterior distribution is then used as an estimate of recombination rate for each core gene in each species.

Using this approach, we inferred the recombination rate of each core gene across all core genomes (*i.e*., the set of genes shared by nearly all the members of a species) of our initial study dataset: 162 bacterial species and one archaeon^20^. Each species was composed of 15 to 100 non-redundant genomes that were classified to a species using both the average nucleotide identity (ANI) and the patterns of gene flow^22^. The core genes whose summary statistics could not be robustly used for inference of recombination rate (*i.e*., that fell outside of the distribution of simulated summary statistics) were excluded from the analysis (see Supp. Methods). We further excluded the species whose core genome was highly reduced (<200 core genes) after excluding those genes. After applying these criteria, our final dataset was composed of a total of 145 bacterial and one archaeal species for which recombination rates of individual core genes could be confidently inferred. We found that the average rate of recombination across core genes within each species was tightly correlated to the species *r/m* previously inferred using the whole core genome (Spearman’s *Rho*=0.91, *P*<10^−15^: Fig. 2)^20^. The number of core genes for which a robust estimate of *r/m* could be inferred varied from a minimum of 218 for *Salmonella enterica* to a maximum of 4,161 for *Burkholderia gladioli.* In total, the *r/m* values were estimated for 208,686 core genes. The data summarizing the size of the core gene dataset, the distribution of *r/m* and the summary statistics for the species included in this analysis are presented Supp. Dataset 1.

**Figure 2.**
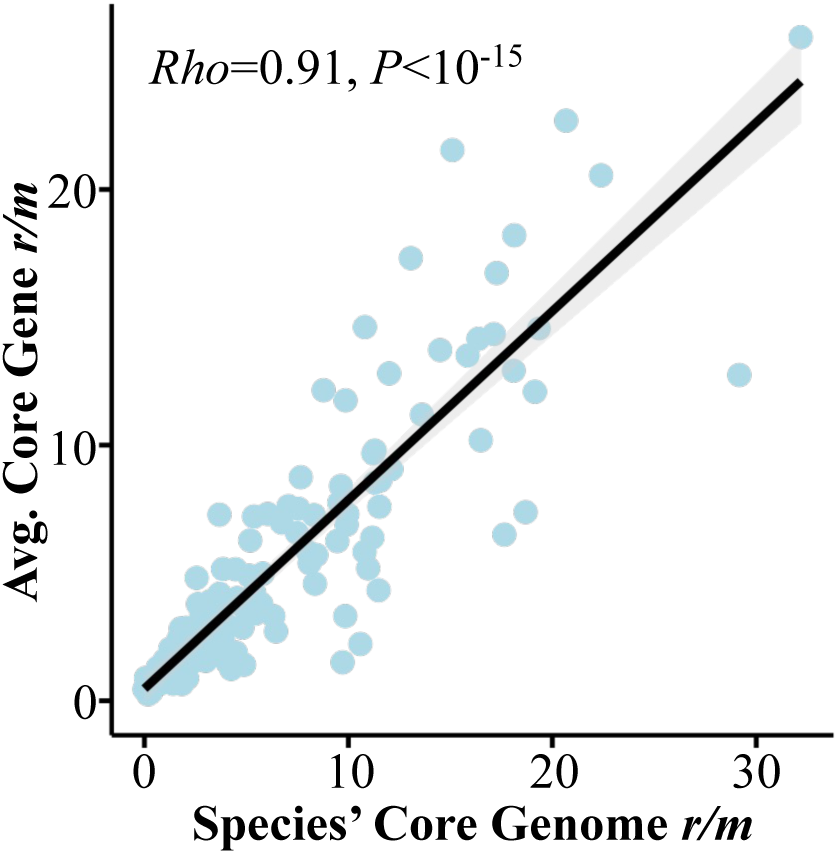
Correlation between core genome *r/m* estimates from Torrance *et al.* (2024)^20^ and the average *r/m* values across core genes for the same species (Spearman’s *Rho*=0.91, *P*<10^-15^).

### Homologous Recombination Rate Variation by Gene Function

We first tested whether rates of homologous recombination vary across gene functional categories (*i.e.,* clusters of orthologous genes (COG)); all genes were classified into COG categories using *eggNOG* (*n*=208,686 core genes)^23,24^. Overall, a significant difference in *r/m* values across COG categories was detected (Kruskal-Wallis, *P*<10^−22^) and recombination rates were then compared for each relevant functional category independently (18 COG categories in total) using a Wilcoxon Test with Benjamini-Hochberg *P*-value adjustment for multiple testing (Supp. Fig 1A). Significantly lower *r/m* values were observed for genes encoding central cellular functions: *i*) cell cycle control, cell division, and chromosome partitioning (COG category D, *P*<10^−5^), *ii*) translation, ribosomal structure, and biogenesis (COG category J, *P*<10^-76^), and *iii*) transcription (COG category K, *P*<0.05). In contrast, significantly higher *r/m* values were observed for genes encoding more diverse functional categories: *i*) energy production and conversion (COG category C, *P*<10^−4^), *ii*) amino acid transport and metabolism (COG category E, *P*=10^−12^), *iii*) carbohydrate transport and metabolism (COG category G, *P*= *P*<10^−5^), *iv*) coenzyme metabolism and transport (COG category H, *P*<10^−3^), *v*) cell wall/membrane/envelope biogenesis (COG category M, *P*<10^−6^), *vii*) inorganic ion transport and metabolism (COG category P, *P*<10^−7^), *viii*) signal transduction mechanisms (COG category T, *P*<10^−3^) and *ix*) defense mechanisms (COG category V, *P*<10^−12^). The same test was performed within species individually (*n*=146). As expected, fewer significant relationships were observed, probably due to the decrease in statistical power. A bar graph showing the number of species which was significant for each COG group is shown in Supp. Fig. 1B.

### Homologous Recombination Rate in Genes Flanking Clusters of Accessory Genes

Accessory genes (*i.e*., non-core genes that are typically found in a single or in a few genomes within a species) are often transferred by Horizontal Gene Transfer (HGT) and the insertion of these sequences can be mediated by homologous recombination or by other mechanisms^25^. In prior studies, the rate of homologous recombination has been inferred to be higher in the regions flanking horizontally transferred accessory genes^12,26,27^. However, to our knowledge only Oliveira *et al*. (2017) analyzed this trend over a diverse set of species. These authors quantified recombination rates by calculating both the number of estimated recombination events and the amount of phylogenetic incongruencies in core genes flanking accessory gene regions^12^. To test whether we observed similar trends in our dataset, we defined accessory gene clusters as blocks of ≥5 consecutive accessory genes and we compared *r/m* estimates of core genes flanking these regions versus *r/m* estimated of the non-flanking core genes (*i.e.* core genes not directly located next to accessory gene regions) for each species (Supp. Dataset 2). We found that the majority of species (*n*=98, 68%) had an increase in recombination rate in flanking core genes relative to non-flanking core genes. However, the increase was only significant in nine species (*Bacillus megaterium, B. wiedmannii, B. cereus, Staphylococcus warneri, S. equorum, Lactobacillus kunkeei, Burkholderia multivorans*, *B. vietnamiensis,* and *Bifidobacterium adolescentis*) (Wilcoxon test, Benjamini-Hochberg adjusted *P*<0.05) and no species were found to have a statistically significant decrease. The increase in *r/m* in the core genes flanking accessory regions was overall relatively small: we measured an average increase in *r/m* across species of 0.47±0.64.

### Rate of Homologous Recombination, GC-content, and Selection

Previous studies have reported a positive relationship between the recombination rate and the GC-content of gene sequences^28–30^. This result has been interpreted as evidence that recombination can enhance genomic GC-content either by *i*) a mechanistic bias during the recombination process (the biased gene conversion hypothesis)^28^ or by *ii*) increasing the effectiveness of selection (the selection hypothesis)^29,30^. We compared our estimates of recombination rates to the average GC-content estimated across all the sequences of each core gene using Spearman’s correlations with Benjamini-Hochberg *P*-value adjustment for all 146 species (Fig. 3A, Supp. Dataset 3). As reported in previous studies, we observed that the relationship between *r/m* and GC-content was positive (*Rho*>0) in most species (*n*=119, 82%).

**Figure 3.**
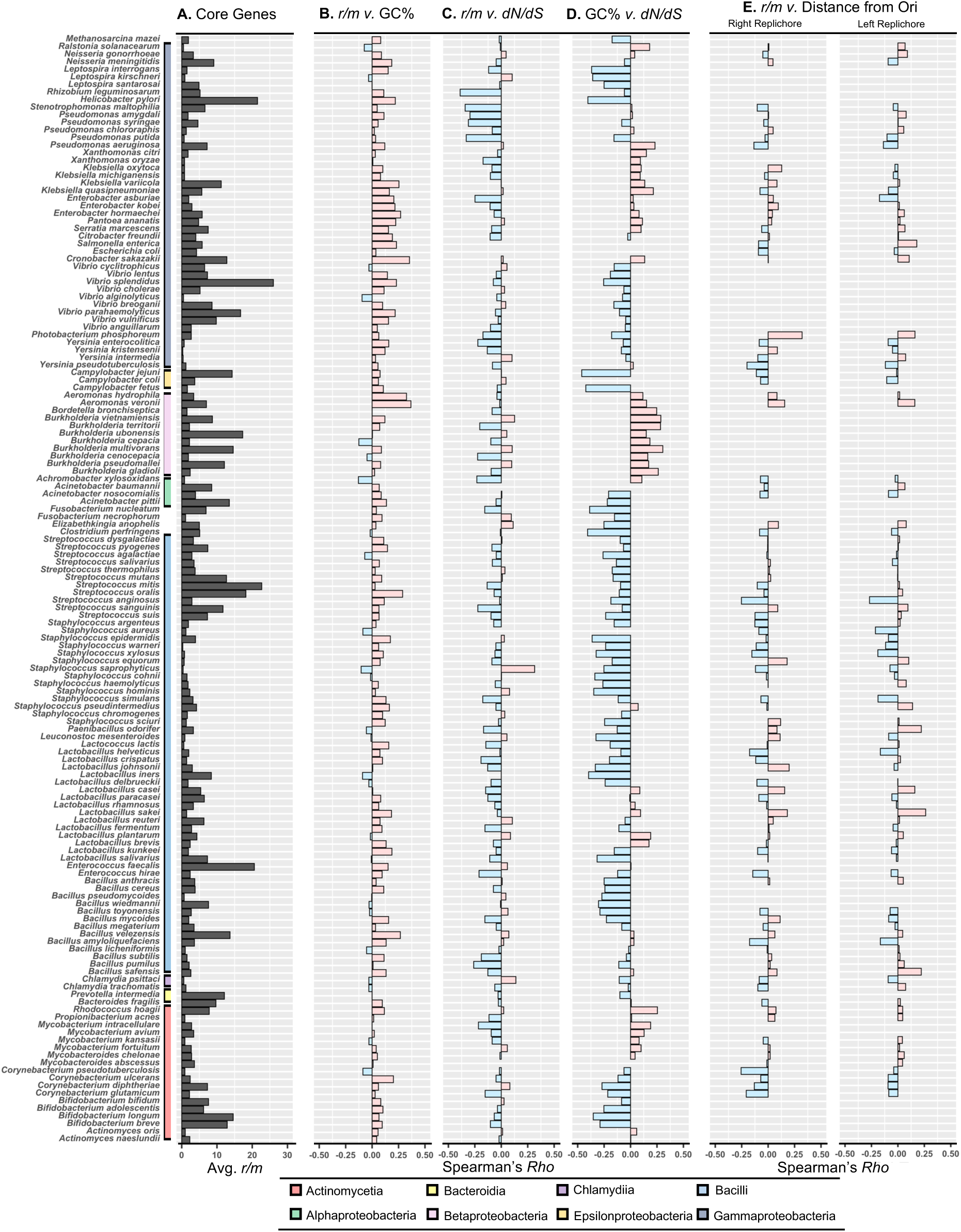
Variation of Spearman’s coefficients of correlations between core gene *r/m vs.* GC content, signatures of selection, and distance from the origin of replication in circular chromosomes. A) A bar graph of the average *r/m* across core genes from each species (*n*=146). Species are organized by phylogenetic class. B) Spearman’s correlation coefficients (*Rho*) are plotted for each species (when available) for *r/m vs.* GC% (*n*=146), *r/m vs. dN/dS* (*n*=142), and GC% *vs. dN/dS* (*n*=142). C) Spearman’s correlation coefficients (*Rho*) are plotted for each species with a circular chromosome (*n*=102, see Supp. Methods) for *r/m vs.* distance from the origin of replication (Ori) for arbitrarily defined “right” and “left” halves of the replichore.

The relationship between *r/m* and GC-content was significant for 83 (57%) of these species, among which nine had a significant negative correlation (6%) and the remaining 74 showed a significant positive correlation (51%).

To investigate whether there is a relationship between *r/m* and the impact of selection, we estimated the ratio of non-synonymous to synonymous substitution rates (*dN/dS*) using PAML for each core gene alignment and for each species (see Methods). Species were included in this analysis if *r/m* and *dN/dS* could be both estimated for at least 200 core genes (*n*=142). We found a significant correlation between *r/m* and *dN/dS* for 73 species (51%) (Spearman’s tests, Benjamini-Hochberg adjusted *P*<0.05) (Fig. 3A). Of those that were significant, 14 showed a positive correlation and the remaining 81% (*n*=59) presented a negative correlation between *r/m* and *dN/dS* (Supp. Fig. 2). These results indicate that genes with higher recombination rates are evolving under stronger or more effective selective pressures. Separately, we observed that *r/m* and *dN* were significantly correlated for 114 species with a positive significant relationship for 67% (*n*=79) and *r/m* vs. *dS* were significantly correlated for 123 species with a positive significant relationship for 80% of species (*Rho*>0, *n*=98). These results further indicate that the correlation between *r/m* and *dN/dS* is not solely driven by *dN* nor by *dS* (Supp. Dataset 4).

We further tested whether a relationship existed between *dN/dS* and GC-content across the core genes of the 142 species (Fig. 3A, Supp. Dataset 5). We observed a significant relationship between *dN/dS* and GC-content across 113 species (77%) (Spearman’s tests, Benjamini-Hochberg adjusted *P*<0.05). Of these, 68% of species had a significant negative relationship between *dN/dS* and GC% (*n*=77, *Rho*<0.0) and the remaining 32% (*n*=36) had a significant positive relationship. It should be noted that the correlations between *dN/dS* and GC% were overall stronger than the correlations between *r/m* and GC% (Fig. 3A). This trend suggests that variations in GC-content along bacterial genomes are primarily driven by selection rather than biased gene conversion. Interestingly, we observed that the strength of the correlation between *dN/dS* and GC-content varied substantially and predictably across species according to their overall GC-content (Spearman’s *Rho*=0.76, *P*<10^−26^, Fig. 4) (Supp. Fig. 3). More specifically, we found that species with overall low GC-content presented stronger negative correlations between genic GC-content and *dN/dS*, suggesting that selection acts more strongly to maintain higher genic GC-content in low GC-content bacteria.

**Figure 4:**
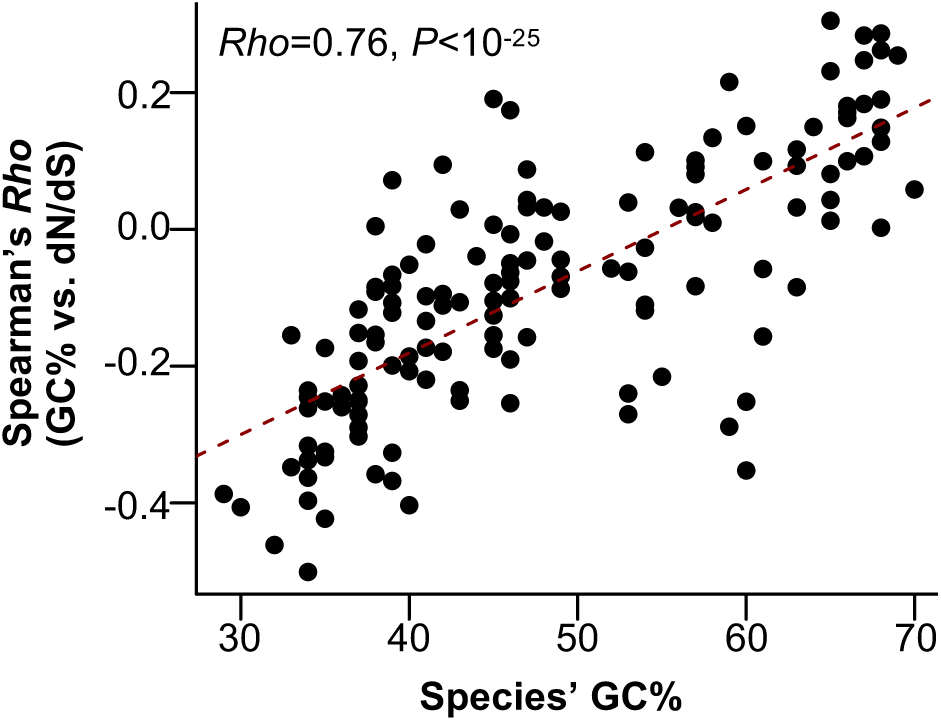
Correlation between GC% vs. *dN/dS* of individual core genes (Spearman’s *Rho*) relative to the overall species’ GC-content (*n*=142) (Spearman’s *Rho*=0.76, P<10^−25^).

### Homologous Recombination Rate and DNA Strand Bias

It has been hypothesized that homologous recombination could be elevated in genes oriented on the lagging strand of replication due to increased frequency of head-on collision between replication and transcriptional machinery and subsequent strand repair mediated by homologous recombination^31^. To test this hypothesis, we compared *r/m* between genes present on the leading and lagging strands using a Wilcoxon test with Benjamini-Hochberg *P*-value adjustment for 102 species with circular chromosomes and where Ori and Ter could be clearly identified (see supplementary methods). We found little evidence supporting a difference in recombination rate between the leading and lagging strands. Of the 102 species assessed, only seven species presented a significant difference in recombination rate across core genes between strands (Supp. Dataset 6). Of those seven, four species had a significant increase in their lagging strand *r/m* (*Pseudomonas chlororaphis*, *Lactobacillus fermentum*, *Staphylococcus saprophyticus*, and *L. kunkeei*) and three species presented the reverse trend (*Yersinia intermedia*, *Serratia marcescens*, and *Lactobacillus salivarus*). Notably, this difference in *r/m* between strands was small (difference in *r/m* average between leading and lagging strands ≤ 1) for all species. Overall, there are no differences in recombination rate between the core genes of the leading strand and the core genes oriented on the lagging strand across bacterial species.

### Evolution of the Genomic Landscape of Homologous Recombination

We next tested whether the patterns of recombination rates were conserved between species following speciation and divergence. We compared the genomic patterns of recombination between all pairs of species within each genus, which represented 109 unique pairs of species. For each pair, recombination rates were compared between shared orthologs to determine whether the patterns of recombination rate were a conserved trait between closely related species (Supp. Dataset 7). We found a positive significant correlation between the recombination rates of the shared orthologs for 36 species pairs (Spearman’s test, Benjamini-Hochberg adjusted *P*<0.05), indicating that genomic landscapes of recombination are somewhat conserved between related species. Conservation in recombination rate was then compared to pairwise divergence between species of the pair. We found that the species with evidence of conservation in recombination rate across shared orthologs had lower pairwise divergence (average A.A. branch length = 0.25±0.18) versus those without evidence of conservation in *r/m* (average A.A. branch length = 0.37±0.30) (Supp. Fig. 4). This result further supports the fact that the genomic patterns of recombination are conserved over short evolutionary distances.

### Genomic Landscapes of Homologous Recombination

Bacterial chromosomes are highly organized entities, and these constrains might shape recombination rates across genomes. We first tested whether recombination rates varied between the origin of replication origin (Ori) and the terminus (Ter) for the 102 species with circular chromosomes for which Ori and Ter could be confidently identified based on the GC-skew (see Methods) (Supp. Dataset 8). We found that *r/m* was higher near Ori in 65% of species (*n*=66) (Fig 3B). Correlations between *r/m* and distance to Ori were statistically significant for 36 species (Spearman’s test, Benjamini-Hochberg adjusted *P* <0.05). The majority of those (*n*=26, 72%) displayed higher rates of recombination near Ori whereas the inverse (*r/m* lower near Ori) was observed and significant in the 10 remaining species (28%). These results indicate that most bacteria present a bias of increased recombination near Ori (Fig. 3B).

We further tested whether recombination rates were symmetrical across both replichores of circular chromosomes. Core gene sets were divided at the *Ori-Ter* axis into “right” and “left” replichores, which were arbitrarily defined. For each replichore, we compared the rate of recombination of the core genes relative to their positions in the *Ori-Ter* axis. Interestingly, we found that the absolute values of Spearman’s *Rho* between both replichores were positively correlated to each other in most species (Spearman’s *Rho*=0.4, *P*<10^−4^) supporting the evidence of some levels of symmetry in the patterns of recombination rates in the two replichores (Fig. 3B, Supp. Fig. 6). This symmetry is also visually apparent in several species where we note the landscape of recombination forms an approximately parabolic graph when plotted linearly from the Ori (Supp. Fig. 6A & B and some species in Fig. 5). Overall, these visualizations support our previous results that the variations of recombination rates along the genomes are roughly symmetrical in both replichores for many species (*i.e*., symmetrical relative to the Ori-Ter axis). Interestingly, although the number of estimable core genes varied across species, we observed conserved genomic landscapes of recombination rates across the species of some genera (*e.g*. *Lactobacillus* (Supp. Fig. 7) and *Staphylococcus* (Fig. 5)).

**Figure 5:**
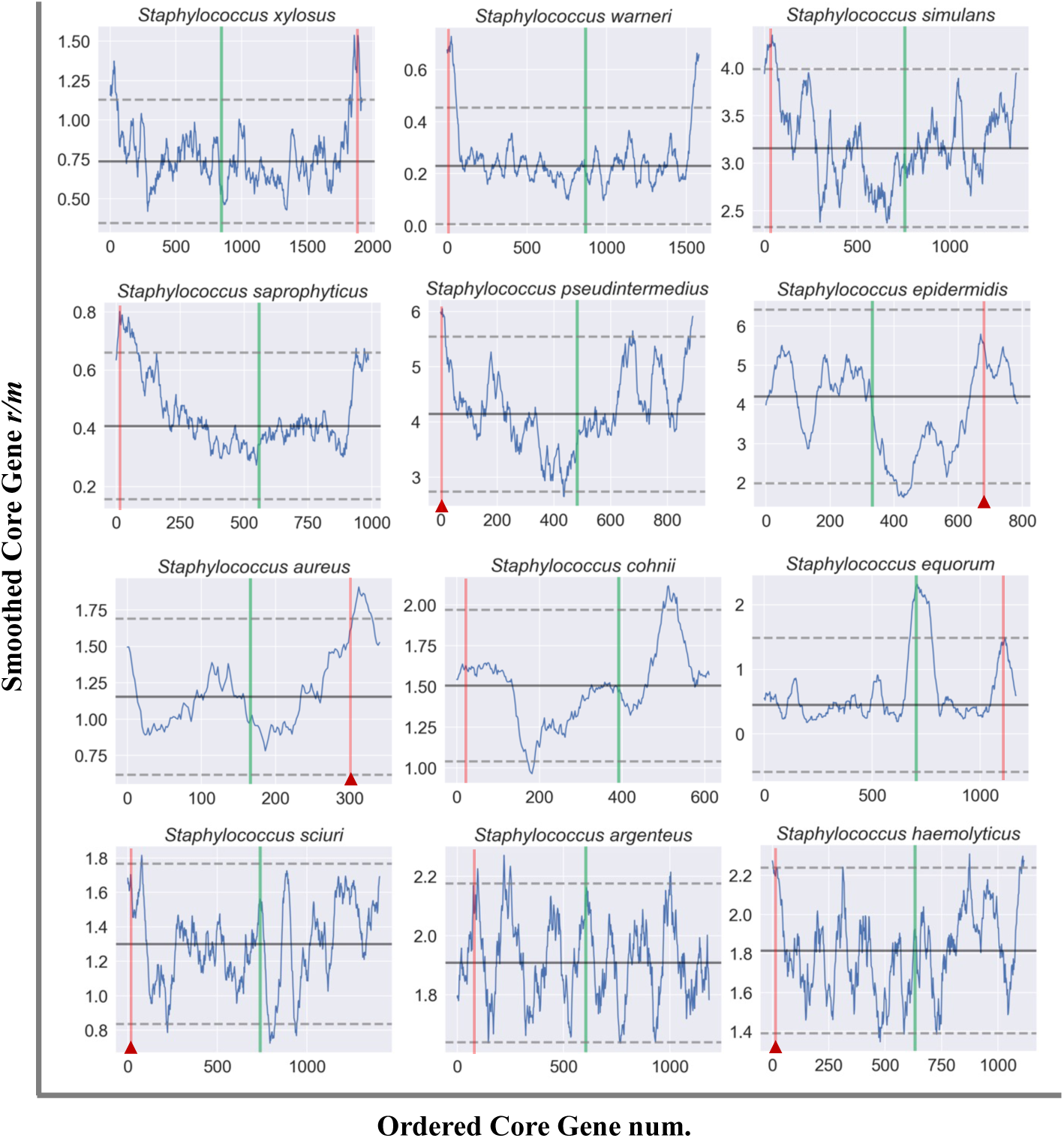
Variation of recombination rate variation along the genomes of *Staphylococcus* species. Each graph represents the smoothed average *r/m* across estimable core genes in a sliding-window of 50 with a step of 2. The location of Ori is at *x*=0 and the location of Ter is demarcated by a green line in each plot. The solid horizontal line represents the mean *r/m* across estimable core genes and the dashed lines represents the *r/m* values two standard-deviations from the mean. The red line on each plot represents the location of *orfX* (a SCC integration site) and the red triangle indicates the presence of SCCmec integration at this site.

Previous studies have found that recombination rates vary extensively across bacterial chromosomes in species such as *Staphylococcus aureus*, *Streptococcus pyogenes*, and *Campylobacter jejuni* with little visible pattern for most species except for several spikes, or “hotspots” of recombination^4,7,8,26^. Hotspots of recombination are of particular interest as they have previously been found to correspond to the presence of genes associated with potentially adaptive traits such as virulence or antibiotic resistance^7,8^. As mentioned above, and in accordance with other studies, we observed that rates of recombination are significantly elevated in genes associated with adaptive traits such as cellular defense among others (see Supp. Fig. 1)^7,10^. To visualize the patterns of recombination along bacterial genomes, we plotted the average of recombination rates across the core genome of each species (*n*=102) using a sliding window (starting at the origin of replication (*x*=0) and with the location of the terminus plotted as a line in green) (Supplementary Folder 1). Sharp local variations in recombination rates were clearly identified in the genome of most species. We defined hotspots and coldspots (regions of particularly low recombination) as regions where local *r/m* differed by more than two standard deviations from the mean across the genome. Overall, we detected many hotspots and coldspots across species with most species having at least one hotspot or coldspot as defined by this metric. In accordance with prior studies, we found that coldspots primarily encode housekeeping genes, whereas hotspots are enriched in genes that are potentially adaptive such as those associated with metabolic or virulence traits^7^.

### Landscape of Homologous Recombination in *Staphylococcus*

We observed that the genomic landscape of recombination was conserved in several genera, but we noted particularly interesting patterns in *Staphylococcus*. For most *Staphylococcus* species (*n*=10, 83%), the genes near Ori were observed to have the highest recombination rate with a gradual decrease moving towards Ter. This trend was found to be statistically significant for at least one replichore in nine species (75%) (Spearman’s test, Benjamini-Hochberg adjusted *P*<0.05). This is a pattern that has, to our knowledge, only been observed previously in *S. aureus*^7,8^. In our study, the shape of the rate of recombination across the genome appears to be conserved across most species within the genus and can be roughly described as an upward parabola (Fig. 5). Interestingly, a gene homologous to a SPOUT-methyltransferase was found within the conserved hotspot proximal to Ori in all species (*n*=12) (Fig. 5: SPOUT-methyltransferase highlighted in red). This finding is notable as this gene, often alternatively referred to as *orfX*, is thought to be the integration site for Staphylococcal Chromosomal Cassettes (SCC) including SCCmec which is a clinically determinant factor used to determine whether a *Staphylococcus* infection is classified as methicillin-resistant^32,33^. Indeed, all but one species contained a region of accessory genes which are likely mobile (*i.e*. encoding putative transposases) adjacent to this gene, indicating that the SPOUT-Methyltransferase homolog is a conserved integration site for a variety of MGEs across *Staphylococcus* species. In fact, MecA (specifically, Penicillin-Binding Protein 2a (PBP2a) family MecA) was annotated within the accessory region proximal to the SPOUT-methyltransferase in the reference genome for five species in our analysis (*S. aureus*, *S. pseudointermedius*, *S. epidermidis*, *S. sciuri*, and *S. haemolyticus*). The fact that this clinically relevant integration site is located within a region with elevated recombination rate indicates that homologous recombination likely contributes to the integration of MGEs in this region and plays a role in transfers of clinically relevant MGE, such as SCCmec.

### Overview of Clinically Relevant Genes in Hotspots of Recombination

Besides *Staphylococcus,* several species had clinically relevant gene sets present in their respective hotspots of recombination. In *B. subtillis*, the hotspot near Ter contains genes annotated as belonging to the *yngABC* operon, which is associated with lipid metabolism, biofilm formation, and anaerobic growth^34^ (Supp. Fig. 8). Several of these genes were found to have *r/m* rates at approximately 10 times the genome average (ex: *yngB r/m*=12.9, *yngI*, *yngL*, and *yngK r/m*>10) contributing heavily to the spike in *r/m* average in a region with an otherwise low recombination rate. The gene *MurM* (*r/m*=9.3, gene 198 in Supp. Fig. 8) is present in a hotspot in *Streptococcus pyogenes* which is linked to penicillin resistance and peptidoglycan formation as well as a regulator of the stringent response pathway in *S. pneumoniae*^35^. The second hotspot of *S. pyogenes* encodes a lactose-specific PTS (phosphotransferase system) (*r/m*∼19, gene 684 in Supp. Fig. 9) which has been found to play a role in Group A *Streptococcus* (GAS) virulence in mice^36^. The single hotspot of *S. mutans* contains genes involved in pyrimidine biosynthesis (*pyrK, pyrD, pyrF, pyrE*: *r/m*∼20), whose upregulation is associated with acid-tolerance in *S. mutans* cultures^37^ and may contribute to their role in generating dental carries. In *Yersinia enterolitica* the main hotspot contains the gene *YadG* (r*/m* =2.7) which encodes a putative ATP-binding protein of an ABC transporter system and was found to be associated with granuloma (*i.e.*, aggregate of host immune cells) formation in *Y. pseudotuberculosis* infection^38^ (Supp. Fig. 10). *Pseudomonas aeruginosa* presents a hotspot of recombination rate in a region encoding the putative PA1272 operon (*Cob(I)alamin adenosyltransferase*) which is involved in Vitamin B12 synthesis and plays a role in bacterial persistence within the host as well as pathogenicity (Supp. Fig. 11)^39,40^. Here, *CobO* has an *r/m*=18.5 and the other associated Cob proteins were found to be accessory genes and so did not have estimable *r/m* rates^41^. Notably, many genes within this hotspot were excluded from the analysis as their summary statistics fell outside of the distribution of simulated summary statistics for *P. aeruginosa* indicating that the true rate of recombination may be underestimated in this region. The hotspot of recombination in *Corynebacterium diptheriae* was associated with ion transport such as the *czcD* gene (*r/m*=14) which is hypothesized to aid in avoidance of macrophage-induced zinc toxicity in human infection^42^ (Supp. Fig. 12). Though not all hotspots of recombination were analyzed for all species in this analysis (*n*=146), all gene annotations and corresponding recombination rate data are available on Kaggle (www.kaggle.com/datasets/ellistorr/bacteria-gene-rm).

## Discussion

Though homologous recombination rate is expected to vary at the genomic scale, its variation across bacterial genomes has, to our knowledge, been examined in only 10 species and often across relatively few genomes and genomic sites^4,7,8,26,43^. In this study, we estimated the recombination rate (*r/m*) for individual core genes (*n*=208,686 total) across 145 bacterial species and one archaeon using an ABC framework^20^. We found that estimates of recombination rate averaged across individual core genes were nearly identical to those inferred for the entire core genome in our previous analysis (Fig. 2) indicating that core gene estimates of *r/m* by this method seem robust. As observed in prior studies, we noted a statistically significant decrease in *r/m* for genes associated with conserved housekeeping functions such as those coding for transcriptional and cellular replication machinery (Supp. Fig. 1A)^7,10^. The inverse was found for genes related to metabolism, signaling, and virulence among others. Though different metrics were used to assign both gene functionality and homologous recombination rate in prior analyses, this was found to be true for virulence associated genes in only three species in^7^ (*Escherichia coli*, *Neisseria meningitidis*, and *S. aureus*) (Supp. Fig. 1A). High recombination rate in^7^ was most frequently found to be associated with genes involved in modulation of the cell surface and we observed similar trends, but also high levels of recombination in genes involved in cellular metabolism and transport.

Among species with a circular chromosomal and where Ori and Ter could be confidently inferred (*n*=102), we found that *r/m* was higher at Ori than Ter for 67% of species. This trend has been observed in previous studies in very few species such as *S. aureus*^7,8^ and it has been experimentally demonstrated in a strain of *Salmonella enterica*^5^. This bias may be due to increased exposure of this region to the recombination machinery during chromosomal replication. Indeed, during replication, Ori-proximal DNA is present in two or more copies relative to Ter-proximal regions, and this likely offers increased opportunities for recombination to occur near Ori. We compared the statistical relationship between the recombination rate of core genes and their distance from Ori separately for each replichore and found that, for many species, the recombination landscape across the two replichores was quite symmetrical (Fig. 3B, Supp. Fig. 6). This trend is particularly interesting because patterns of symmetry are difficult to detect, especially since we could only infer the recombination rate of the core genes and not the entire genome. Studies have found the mutational patterns in bacteria are also symmetrical in both replichores and vary with replication timing^14,44^. Thus, it is likely that replication timing and stalling similarly impacts the variation in homologous recombination rates as well across the bacterial chromosome.

Interestingly, in some genera (*e.g.*, *Staphylococcus* (Fig. 5) and *Lactobacillus* (Supp. Fig. 7)) several species displayed similar landscapes of recombination. A previous study found evidence that recombination rate may be conserved across orthologs of closely related species (*n*=3 species pairs^7^) which may contribute to some of the similarities we observed in *r/m* across genomic landscapes. Across 109 species pairs (within the same genus) we found that 36 had statistically significant correlation in *r/m* across shared orthologs (Supp. Fig. 5). We found that species pairs with conserved recombination landscapes (*n*=36) were more related to one another by comparing the pairwise divergence between significant and non-significant groups (Supp. Fig. 4). Thus, our results provide further evidence that recombination rate across shared orthologs is conserved across closely related species. It is unclear, however, whether this trend is driven by a conserved architecture of the genomes of these species and an overall conservation of the recombination landscape or whether homologous recombination rate is similar in syntenic genes with similar functions and similar selective pressures.

Increased substitution rate has been observed in the genes of the lagging strand of DNA relative to those on the leading strand and some authors have suggested that this may be the result of asymmetrical mutation rate^45^, whereas others have shown evidence that this is due to the higher prevalence of genes evolving under stronger purifying selective pressures on the leading strand^46^. Head-on collisions between the replication machinery and the transcription apparatus are thought to be more common on the lagging rather than the leading strand, and this is hypothesized to have a detrimental impact on gene expression and possibly replication^46^. In contrast to these asymmetrical patterns of substitution rates, we observed a statistical difference in recombination rates across genes present in the leading strand relative to those found on the lagging strand for only seven species (Supp. 6). Overall, it indicates that the asymmetry of the DNA strands does not impact the rates of homologous recombination.

Though the base composition of the genome varies across prokaryotes, the GC-content across the genome is typically conserved at the genus and phylum level^47^. Variations in GC-content have been proposed to be shaped by selective forces (*e.g*., for transcription or translation efficiency) or neutral processes such as mutational biases or biased gene conversion, or a combination of both^47^. Specifically, GC-biased gene conversion (gBGC) is a force shaping the base composition of Eukaryotic genomes whereby mismatches introduced during recombination events are preferentially repaired intro G’s and C’s rather than A’s and T’s. Therefore, in Eukaryotes, regions of high GC-content also display elevated recombination^48^. GC-biased gene conversion is expected to be a neutral process that perhaps counteracts the mutational spectrum that is universally biased towards A and T^49^. However, whether gBGC plays a role in shaping the base composition of Prokaryotic genomes is subject to debate because higher recombination rates are also expected to disrupt genome linkage and thereby enhance the effectiveness of selection^28,47,50^. In accordance with a previous analysis^7^, we observed a positive correlation between *r/m* and GC-content across most species (Fig. 3) which is expected under the gBGC model and the selective model. However, we found a negative correlation between *r/m* and signatures of selection (*dN/dS*) (Fig. 3, Supp. Dataset 4) in most species, and *dN/dS* and GC-content were also correlated (Supp. Dataset 5). Interestingly, we observed that this correlation between *dN/dS* and GC-content was strongly dependent on the overall GC-content of the species. Most notably, species with low GC-content presented a much stronger correlation between genic GC-content and the strength of purifying selection. It likely reflects the enhanced selective pressures acting on the species with low GC-content to maintain G’s and C’s in their genes. These results strongly suggest that GC-content is driven by selective forces rather than gBGC in bacteria.

The presence of recombination hotspots (*i.e*., regions with high relative recombination rate) across the genomes of bacteria is thought to be associated with adaptive genes, including those associated with virulence and pathogenicity because these genes frequently arise from horizontal transmission. Since the transfer and integration of these elements is expected to be partially mediated by homologous recombination, we expected to observe higher recombination rates, or hotspots, in some of these regions^4,43^. We observed that the number and frequency of hotspots varied extensively across bacterial chromosomes of different species as was previously observed for ten species^7^. Although few of these genes within these regions have been described in previous analyses, we did observe that *ksgA* in *S. pyogenes* was similarly elevated in both^7^ and our analysis. Furthermore, in the genus *Staphylococcus*, we observed a strong trend of a decreasing *r/m* with increasing distance from Ori and hotspots of *r/m* proximal to Ori, and this had been observed in two prior studies in *S. aureus*^7,8^. Here, the authors of ^7,8^ noted the presence of the gene *OrfX* (a SPOUT-methyltransferase homolog) in recombination hotspots which acts as an integration site for clinically relevant mobile genetic elements (MGEs) such as SCCmec (a Staphylococcal Cassette Chromosome carrying genes implicated in methicillin resistance).

Interestingly, we found that this integration site is always associated with the recombination hotspot proximal to Ori, which is conserved in all 12 *Staphylococcus* species in our analysis (Fig. 5; *OrfX* highlighted in red). Furthermore, a MGE was found to be integrated at this site in all but one species, and an SCCmec-like element (*i.e.,* an MGE encoding a *MecA* (*PBP2a*) gene specifically associated with high-level of methicillin resistance^51^) was found present at this site in five species (Fig. 5; red triangle denotes presence of SCCmec). Due to the relative conservation of *OrfX* in this highly recombining region proximal to Ori across *Staphylococcus* species, it is likely that an MGE such as SCCmec could be efficiently transferred between species in this genus. Thus, it is probable that all *Staphylococcus* species have the capacity to acquire methicillin resistance through inter-species transfers of elements such as SCCmec. Furthermore, this indicates that a hotspot of recombination, which is highly adaptive, has been conserved for a long period of time.

Finally, it is expected that homologous recombination plays some role in the acquisition of accessory genes through horizontal gene transfer^12^. In fact, prior studies have estimated homologous recombination rates to be elevated in regions flanking MGEs^12,27^. These mobile regions are associated with tracts of accessory genes and so, in this analysis, we compared recombination rates of core genes flanking clusters of accessory genes to all other core genes. Though we found that flanking core genes had a slight elevation in recombination rate across all species, we observed this elevation to be statistically significant in only nine. However, the differences between studies may be due to differences in defining accessory regions or recombination rate. Perhaps as well, these results should be nuanced because many mobile elements do not rely on homologous recombination for integration. For instance, many mobile elements like most temperate bacteriophages encode their own site-specific integrase, and these enzymes are not expected to leave a signal of homologous recombination.

Overall, our results show that homologous recombination plays a major role in shaping the architecture and the evolution of bacterial genomes. Recombination rate is highly variable across the bacterial genome and varies with both gene functional roles and chromosomal structure. Thus, homologous recombination represents a force that substantially contributes to genome plasticity in Prokaryotes. One limitation of our methodology is that we were not able to estimate recombination rates for all genes because some displayed parameters that fell outside of the distributions of the simulated range (Supp. Dataset 1). Thus, the number of hotspots of recombination may be underestimated for some species, and methodological improvements may yield a more complete picture of homologous recombination variation across Prokaryotic genomes. Nevertheless, our results provide a rather complete picture showing how recombination rate is associated with many aspects of the architecture and evolution of bacterial genomes: gene function, replication timing, the origin and terminus of replication, GC-content, selection, and gene transfers.

## Supplementary Material

### Methods

#### Data assembly

The dataset of bacterial species used in this analysis was derived from Torrance *et al.* (2024)^20^ and will be described briefly in this section. Species were included when they presented ≥15 sequenced non-identical genomes available on RefSeq. The assembly quality of the genomes within each species was then ensured by checking that each genome contained the expected number of universal genes as in Raymann, *et al.* (2015)^52^. Further, the species’ borders were also verified and redefined as in Diop *et al*. (2022) by ensuring the genomes adhered to their given species definition by both *i)* ANI (≥ 94% pairwise identity across the core genome) and *ii)* gene flow analysis (*i.e.* genomes were only included in a species if they were inferred to be engaging in gene flow with other members of the species)^22^. For the 162 bacterial species and one archaeon that contained ≥15 species, orthologs were inferred using *CoreCruncher* with default parameters and orthologous genes were defined as core genes when present in at least 90% of the genomes of a species^53^. The core genes were aligned using Mafft^54^. One reference genome was chosen for each species by having, first, the most complete genome assembly (i.e., the least number of contigs) and second, the highest number of predicted coding sequences. Accessory genes were defined as all the other genes that were not defined as core present in the reference genome of each species. A core genome phylogeny was generated from the core gene concatenate for each species using *RAxML* v8 with a GTR+gamma model^55^.

#### Estimation of Gene Recombination Rates

A set of 500,000 forward-in-time simulations with varied rates of homologous recombination (*r/m*) was generated using each species tree and each core genome concatenate using *CoreSimul* through the *recABC* pipeline as in Torrance *et al.,* (2024)^20^. For each species, the recombination rate of each core gene was estimated by comparing the signature of recombination between each core gene alignment and the simulated sequences using ABC. The recombination rate of each core gene was estimated by the recombination rate used to evolve the simulated sequences that most closely and robustly matched the signatures of recombination to the gene. Here, the summary statistics *π* (i.e., average pairwise polymorphisms) and *h/m* (*i.e.* the ratio of homoplasic to nonhomoplasic alleles) were calculated for each core gene within each species. Then, using ABC, the summary statistics for each core gene were compared to the summary statistics generated from each simulated species population with a tolerance of 0.01% to determine the most probable rate of recombination for each gene.

As expected, not all summary statistics of all genes fell within the distribution of simulated summary statistics of the simulated sequences. These genes were removed from further analysis when the summary statistic for a given gene varied from the average of its nearest simulated sequence set by more than ±0.1 for *h/m or* ±0.01 for *π*. This threshold was determined by comparing the graphs and correlations (*Spearman’s rho*) of each simulated summary statistic *vs*. real summary statistic (*h/m* and *π*) for each species. Here, we found that the removal of genes outside these tolerance levels yielded the highest number of robust gene estimates where simulated summary statistics were very close to real summary statistics was (*x=y*, or *Spearman’s Rho* ≥0.97) (see figures in Supplementary Folder 2). The number of genes that were excluded from further analysis by this metric are listed in Supplementary Dataset 1.

Furthermore, some core genes had no polymorphisms at all and thus *r/m* could not be estimated (*i.e., m*=0) and those genes were also excluded from the analysis. These genes are denoted as a *r/m* of “NA” in column 2 of the datasets available on Kaggle (www.kaggle.com/datasets/ellistorr/bacteria-gene-rm). Species with fewer than 200 inferred core genes for which *r/m* could be predicted were excluded from further analysis (*n*=10). Thus, our final dataset was composed of 146 species. The species included in this analysis and the total number of core genes and accessory genes, as well as the number of core genes which had no polymorphisms or had summary statistics which fell outside of the range of simulated summary statistics are detailed in Supplementary Dataset 1. The average rate of recombination estimated across all genes was compared to the *r/m* values estimated on the core genome concatenate generated in Torrance et al. (2024)^20^ in Fig. 2. Detailed data for each species describing the summary statistics, *r/m* estimates of each core gene, and other gene metrics and features amassed in this study is available on Kaggle (www.kaggle.com/datasets/ellistorr/bacteria-gene-rm).

#### Identification of Ori and Ter

The origin (Ori) and terminus (Ter) of replication were identified using the cumulative GC-skew (CGC-skew) in the reference genome for each species as in^56^. We built a graph for each species by plotting the CGC-skew using a 10kb sliding window. Species were only included in this analysis if they had a single clear maximum (corresponding to the approximate location Ter) and a single clear minimum (corresponding to the approximate location of Ori) (*n*=110 species)^56^. In other words, Ori and Ter were not inferred for the species presenting additional local maxima or minima. Species with known linear chromosomes or multiple chromosomes were identified from^57^ and excluded from the analyses of recombination symmetry. Using the 102 species with a single circular chromosome and clearly defined Ori and Ter locations, the recombination rate of each core gene was plotted relative to the absolute distance from Ori and Ter to determine whether recombination rate varied on both replichores of each species. These species were also used to compare *r/m* variations between leading and lagging strand genes. Leading and lagging strands were also defined using the GC-skew where core genes present on the strand with increasing GC-skew with increasing distance from the Ori and a positive orientation were determined to be leading strand genes while those with a negative orientation in this region were determining to be lagging strand genes. In the segment of chromosome with a decreasing GC-skew with increasing distance from the Ori, the positively oriented genes were inferred to be located on the lagging strand whereas the negatively oriented genes were inferred to be located on the leading strand. Variations of recombination rate along the core genome of each species were computed using a sliding window of 50 core genes and a step of two genes. The plots are ordered by using Ori as the starting coordinate (Supplementary Folder 1). Hotspots and coldspots of recombination were defined as genome locations where *r/m* differed by more than two standard deviations from the average *r/m* of the species.

#### Other Gene Analyses

To determine whether recombination rate varied significantly across gene functions, the reference genome of all species (*n*=146) was annotated using *EggNOG*^23^ and recombination rate was compared across 18 COG (clusters of orthologous genes) categories (208,686 core genes)^24^. For this analysis, the COG categories A (RNA processing and modification) and B (chromatin structure and dynamics) were excluded because these categories correspond to Eukaryotic gene functions. Category S was excluded because it corresponds to groups of orthologs without known functions. Furthermore, genes were only compared if they were annotated as belonging to a single COG category and genes that were not assigned to a category were excluded. For each species, a Wilcoxon test with Benjamini-Hochberg *P*-value adjustment was conducted where the recombination rates for the genes in each COG category were compared to the recombination rates of the genes classified in all the other COG categories. Additionally, we also conducted this test while pooling all genes of all species together. We used the same dataset of 146 species to determine whether *r/m* varied with any appreciable pattern in genes flanking accessory gene clusters to those not flanking accessory regions. To do this, we defined accessory regions as regions of the genome containing at least five consecutive accessory genes. We then compared the recombination rate of the core genes flanking these regions to the recombination rate of non-flanking core genes using a Wilcoxon test with Benjamini-Hochberg adjustment for each species.

## Supplementary Figures

**Supplementary Figure 1.**
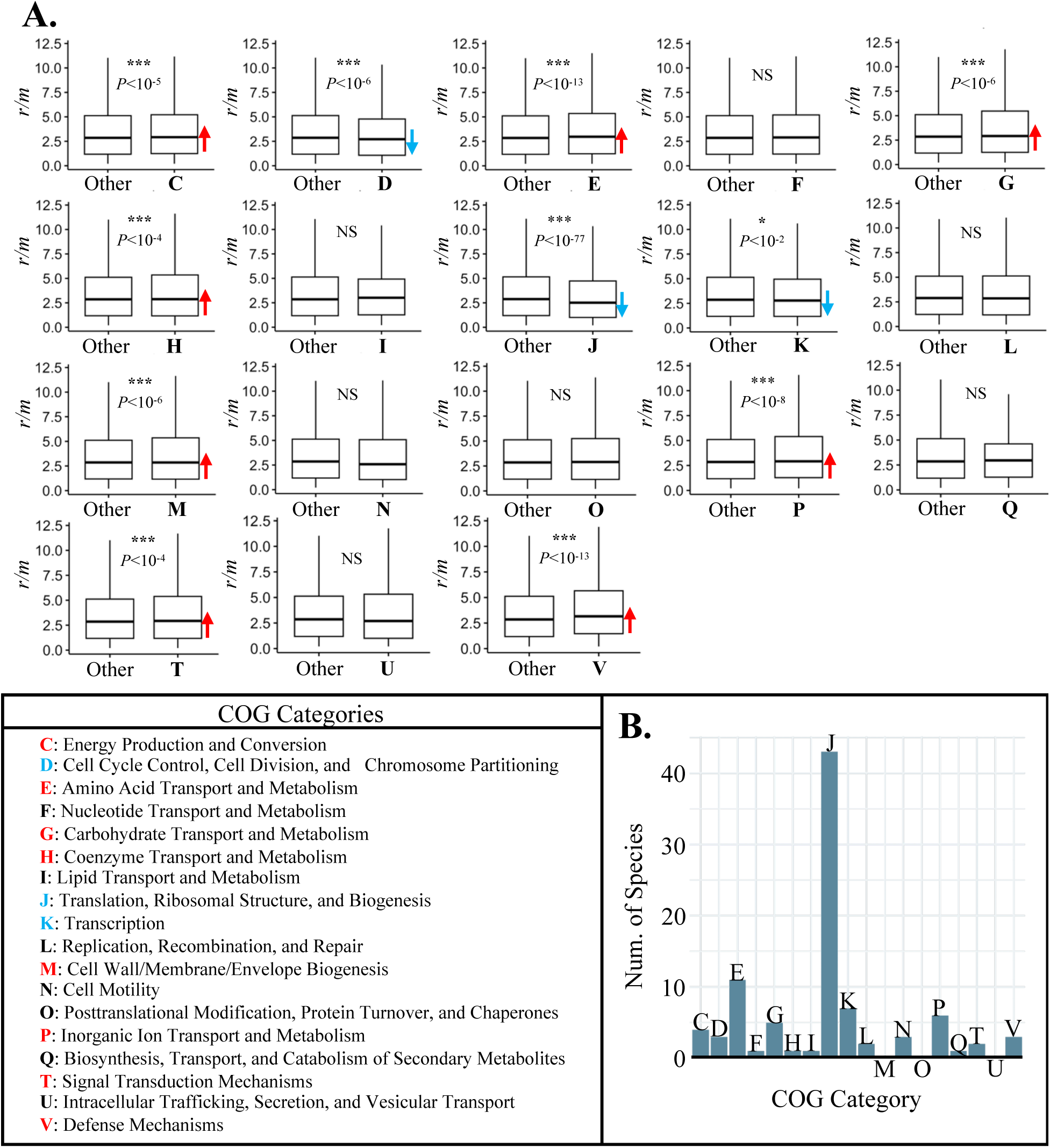
Variation in *r/m* across gene functional categories. A) Boxplots show the variation in *r/m* between each COG (Conserved Ortholog Group^24^) category and all other genes (“Other”). On each plot, significance and Benjamini-Hochberg adjusted *P*-value for each Wilcoxon test is indicated. Red (higher *r/m*) and blue (lower *r/m*) arrows are shown for each COG category that had a significant difference with all other genes. A key listing the description for each COG category is shown in the bottom left. B) A bar graph shows the number of species which had a significant difference in *r/m* between the COG category and all other genes (Wilcoxon test, Benjamini-Hochberg adjusted *P*-value<0.05).

**Supplementary Figure 2.**
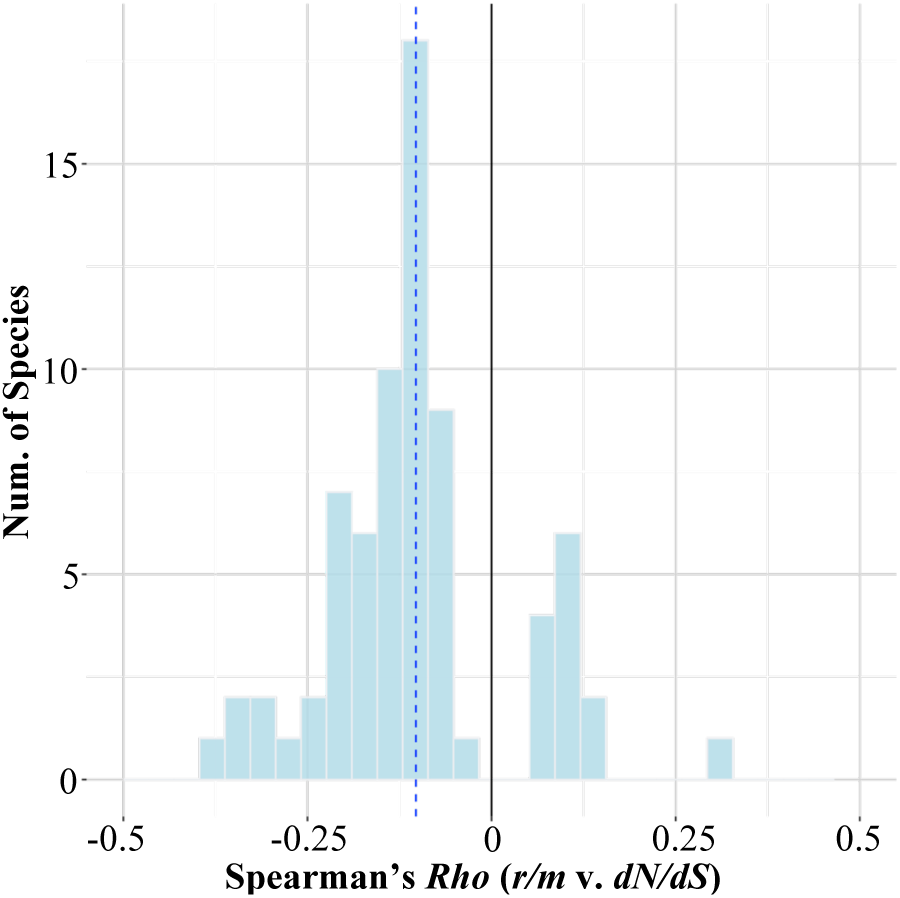
Histogram of significant Spearman’s *Rho* values (Benjamini-Hochberg adjusted *P*<0.05, *n*=73) for the correlation between *r/m* value and *dN/dS* estimated across species’ core genes. The blue dashed line shows the average *Rho* value (Average *Rho*=-0.10**±**0.13).

**Supplementary Figure 3.**
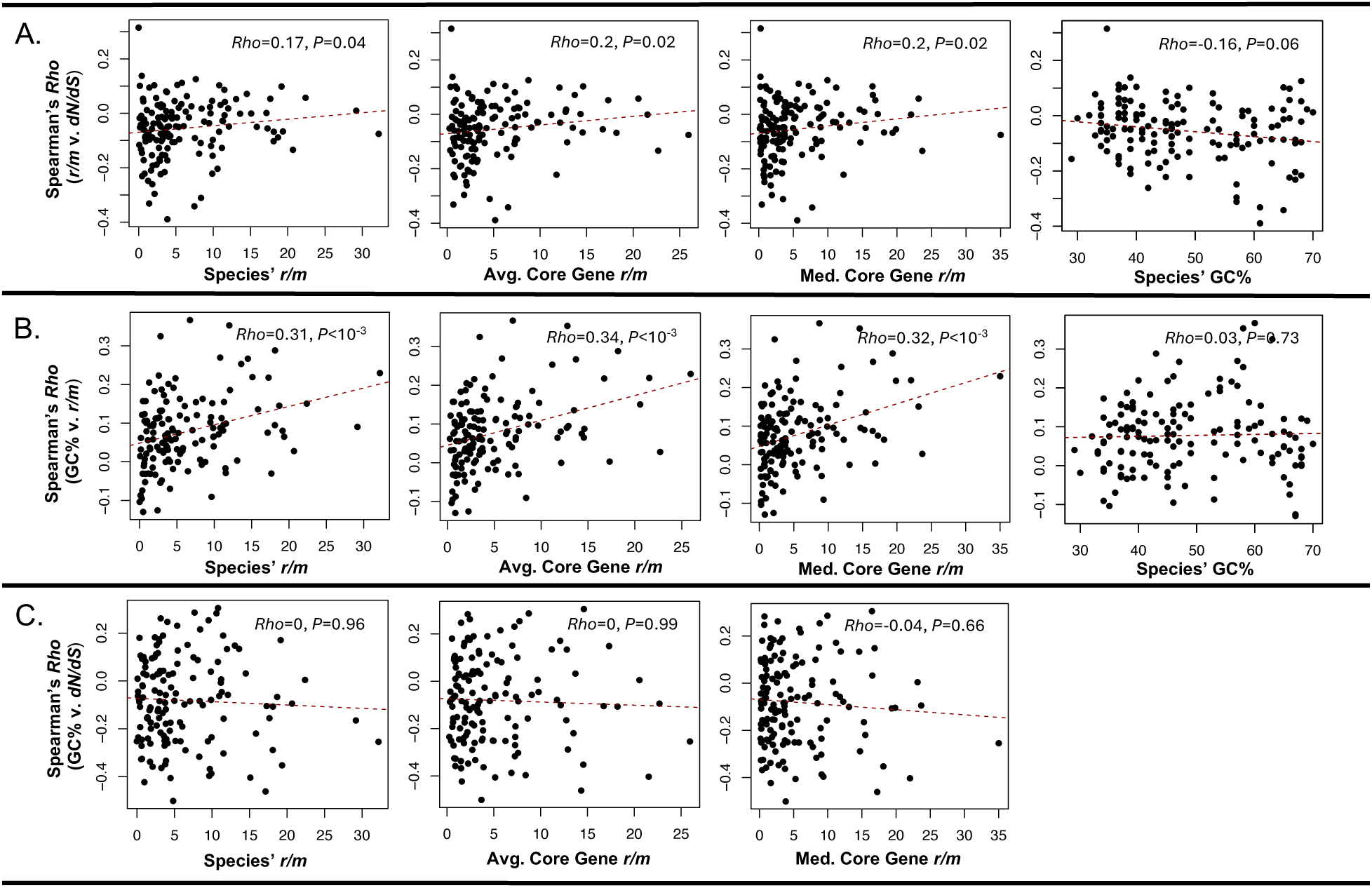
A) Relationship between Spearman’s *Rho* (between core genes’ *r/m* vs. *dN/dS*) and other variables: genome-wide estimated of *r/m* from Torrance *et al.* (2024), average core gene *r/m*, median core gene *r/m*, and species’ GC%. B) Relationship between Spearman’s *Rho* (between core genes’ GC% vs. *r/m*) and other variables: genome-wide estimated of *r/m* from Torrance *et al.* (2024), average core gene *r/m*, median core gene *r/m*, and species’ GC%. C) Relationship between Spearman’s *Rho* (between core genes’ GC% vs. *dN/dS*) and other variables: genome-wide estimated of *r/m* from Torrance *et al.* (2024), average core gene *r/m*, and median core gene *r/m*. The comparison between *Rho* and species GC% is shown in Fig. 4 of the main text.

**Supplementary Figure 4.**
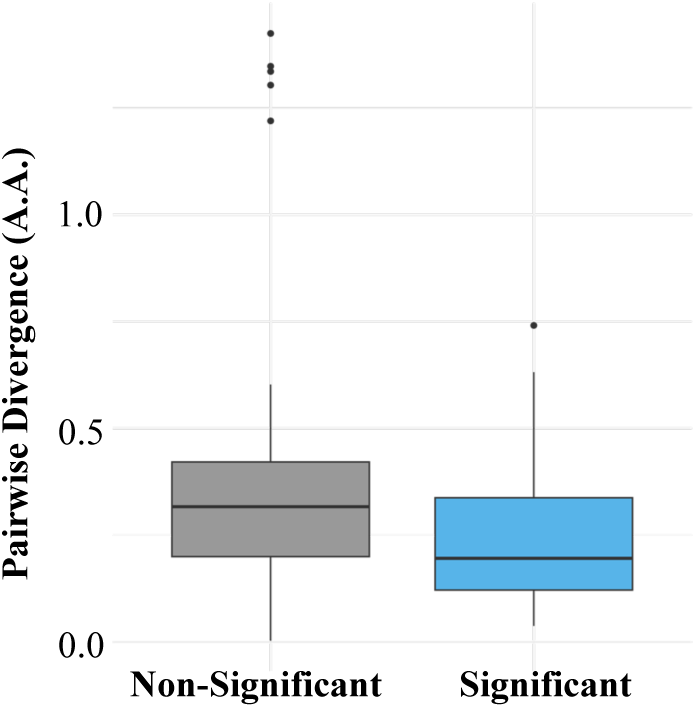
Boxplot comparison of the pairwise divergence (i.e., average branch length: amino acid substitutions per site) for species pairs (*n*=109 species pairs) which had significant correlation in *r/m* values across shared orthologs (“Significant”, *n*=36 species pairs) and those that did not (“Non-Significant”, *n*=73). Only species that belonged to the same genus were compared to one another.

**Supplementary Figure 5.**
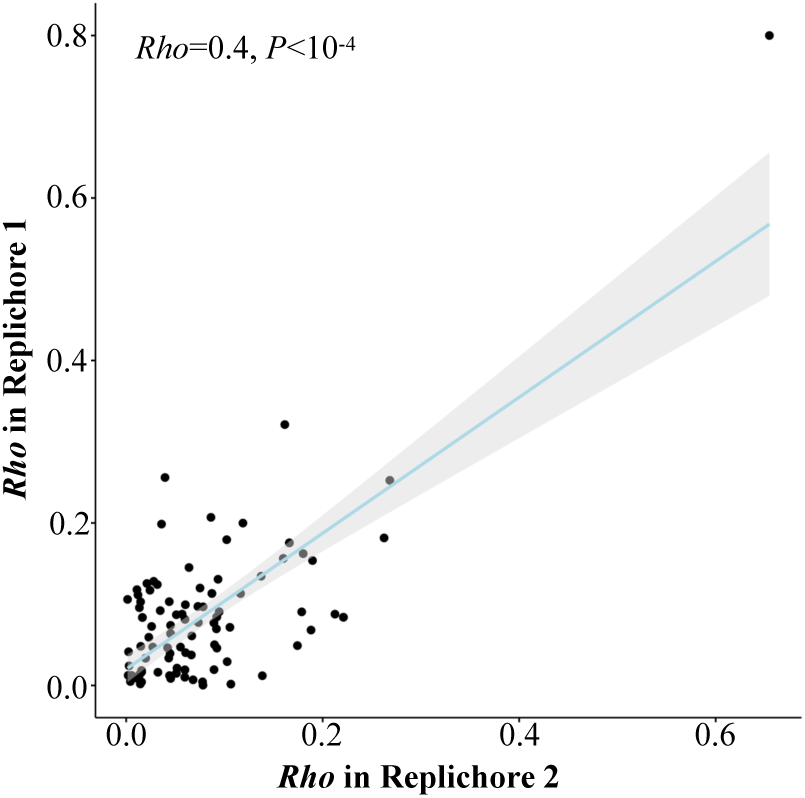
Correlation between Spearman’s *Rho* coefficients from the correlation comparing *r/m* to the distance from Ori of the core genes in the “left” replichore and in the “right” replichore for *n*=102 species with circular chromosomes. Left and right replichores were assigned arbitrarily.

**Supplementary Figure 6.**
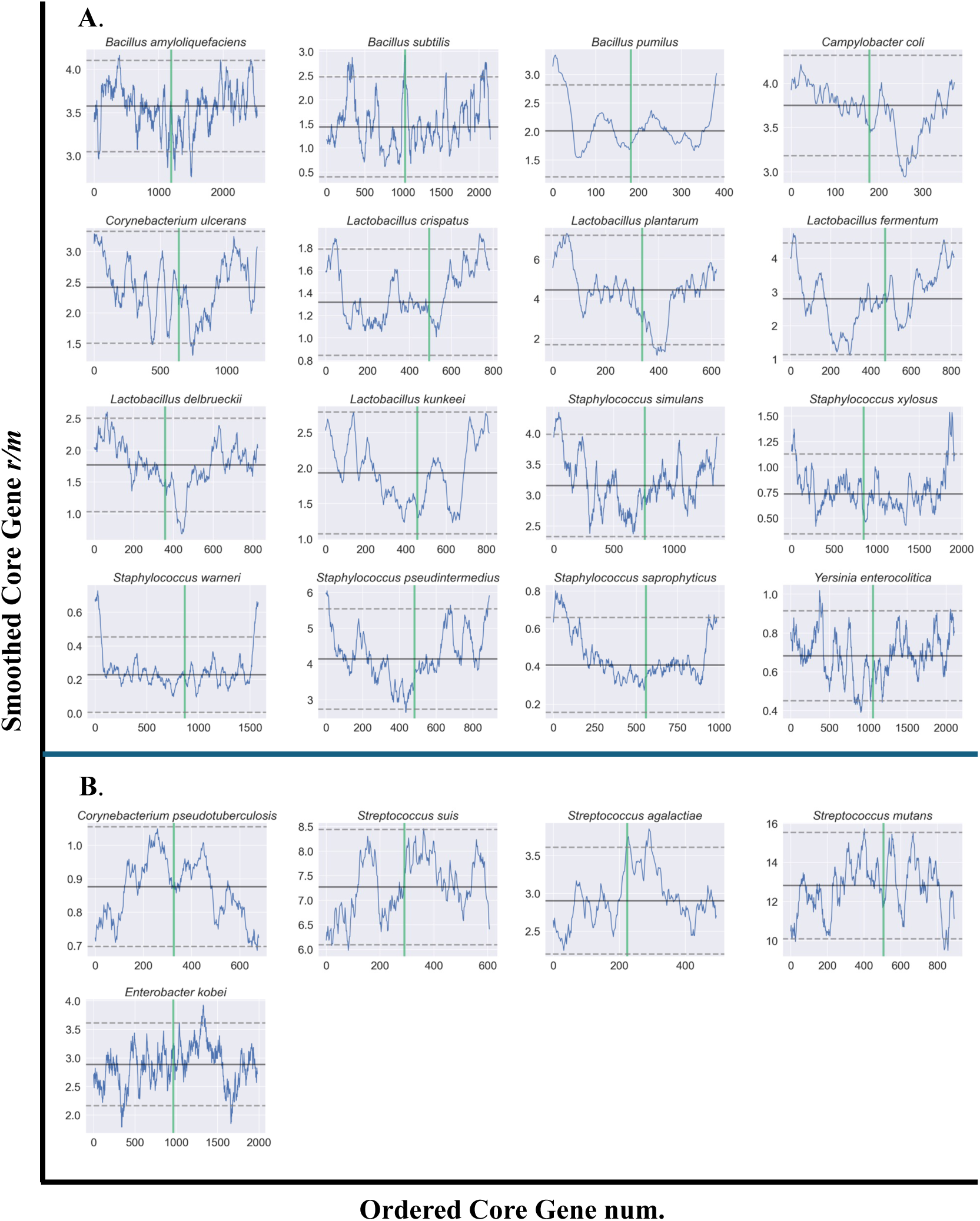
Landscape of recombination rate across bacterial genomes. A) Some bacterial species (*n*=16) displayed an upward “parabola-like” shape of recombination rate variation across their genomes. B) Some bacterial species (*n*=5) displayed a downward “parabola-like” shape of recombination rate variation across their genomes. These graphs display the smoothed average of a sliding window of 50 estimable core genes with a step of 2 where *x* is the core gene number in order of its appearance with increasing distance from Ori. For each graph, *x*=0 corresponds to Ori and the green line demarcates Ter. The black horizontal line denotes the average *r/m* value of all estimable core genes, and the dashed lines represent two standard-deviations from the mean *r/m* in both directions.

**Supplementary Figure 7.**
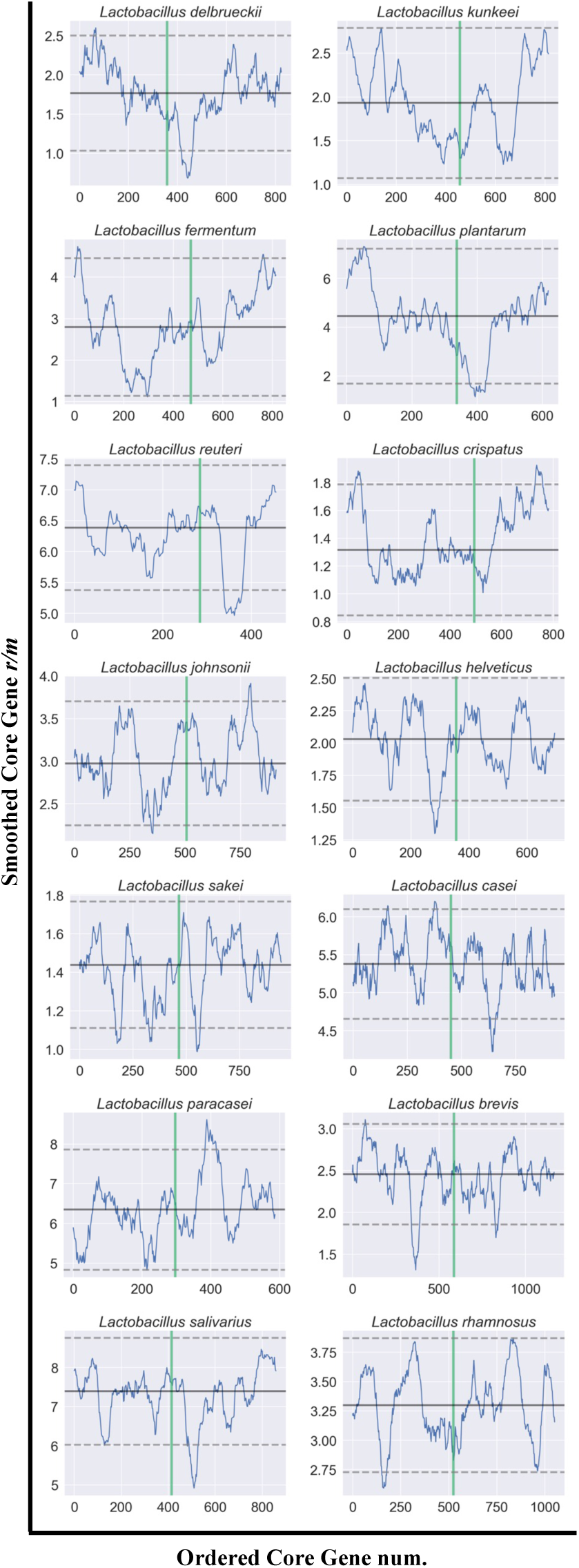
Landscape of recombination rate across *Lactobacillus* genomes (*n*=14). The graphs display the smoothed average across 50 estimable core genes with a step of 2 where *x* is the core gene number in order of their appearance with increasing distance from Ori. For each graph, *x*=0 corresponds to Ori and the green line demarcates Ter. The black horizontal line denotes the average *r/m* value of all estimable core genes, and the dashed lines represent two standard-deviations from the mean *r/m*.

**Supplementary Figure 8.**
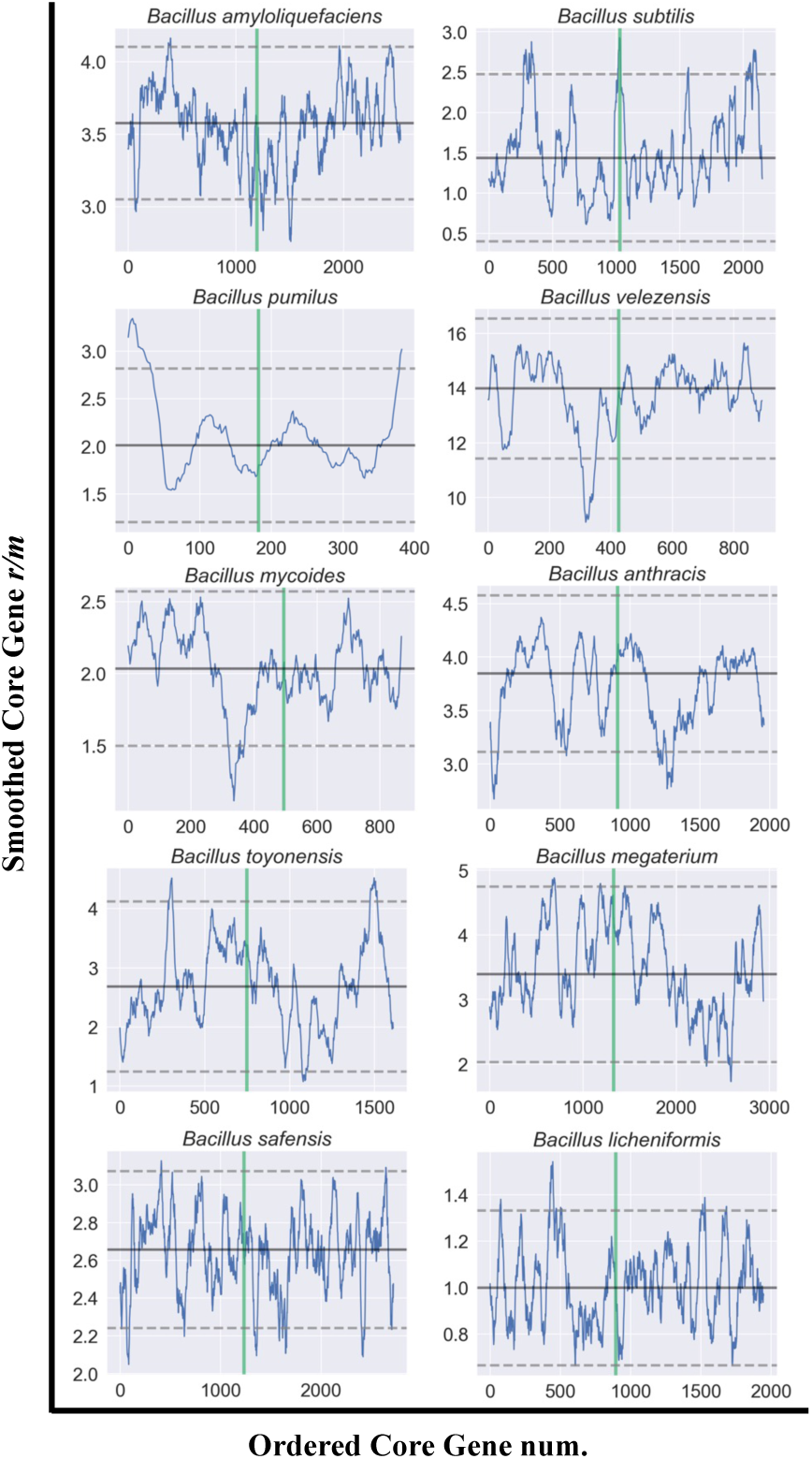
Landscape of recombination rate across *Bacillus* genomes (*n*=10). These graphs display the smoothed average across 50 estimable core genes with a step of 2 where *x* is the core gene number in order of its appearance with increasing distance from Ori. For each graph, *x*=0 demarcates the Ori and the green line demarcates the Ter. The black horizontal line denotes the average *r/m* value of all estimable core genes, and the dashed lines represent two standard-deviations from the mean *r/m*.

**Supplementary Figure 9.**
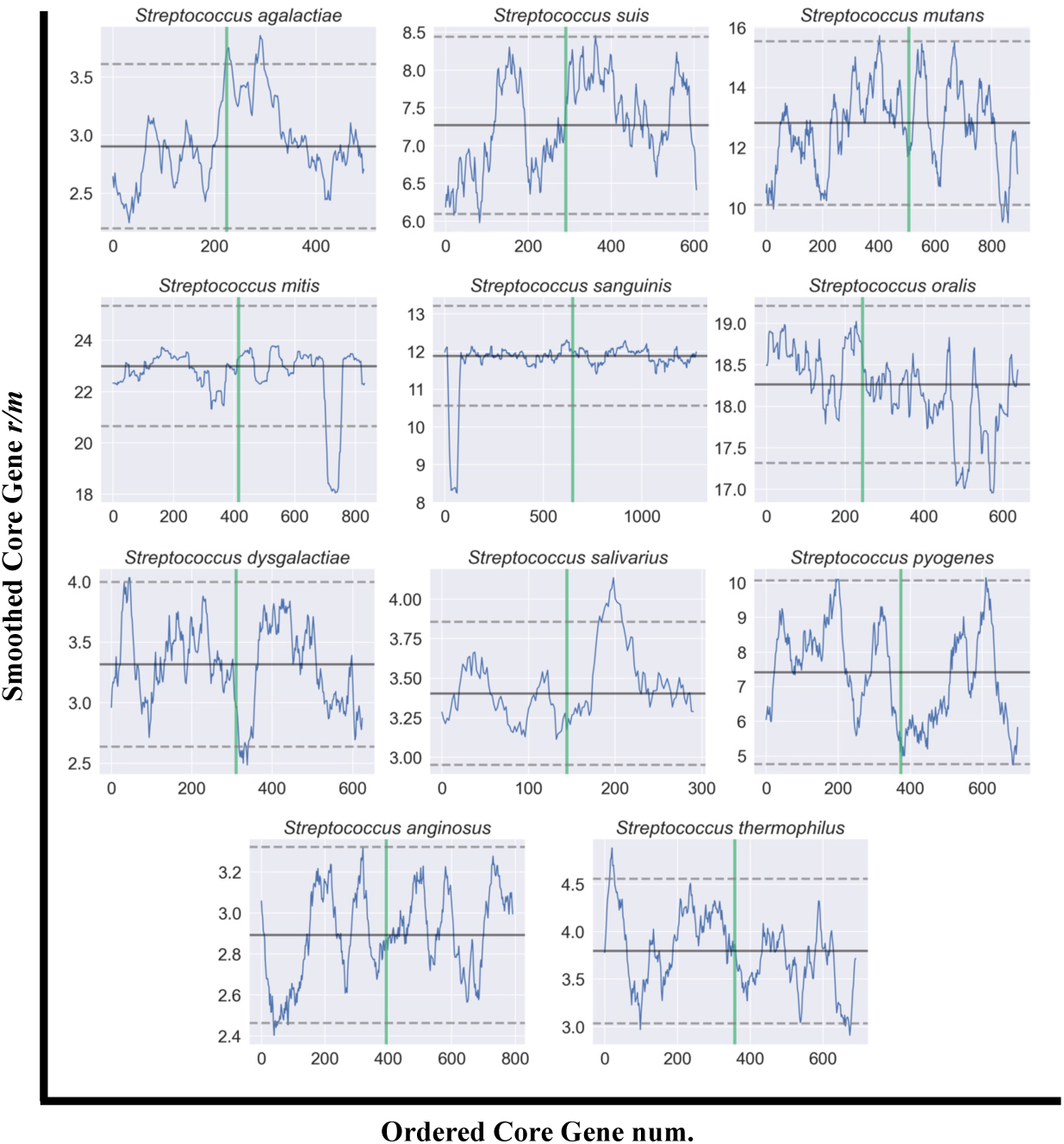
Landscape of recombination rate across *Streptococcus* genomes (*n*=11). These graphs display the smoothed average across 50 estimable core genes with a step of 2 where *x* is the core gene number in order of its appearance with increasing distance from Ori. For each graph, *x*=0 demarcates Ori and the green line demarcates Ter. The black horizontal line denotes the average *r/m* value of all estimable core genes, and the dashed lines represent two standard-deviations from the mean *r/m*.

**Supplementary Figure 10.**
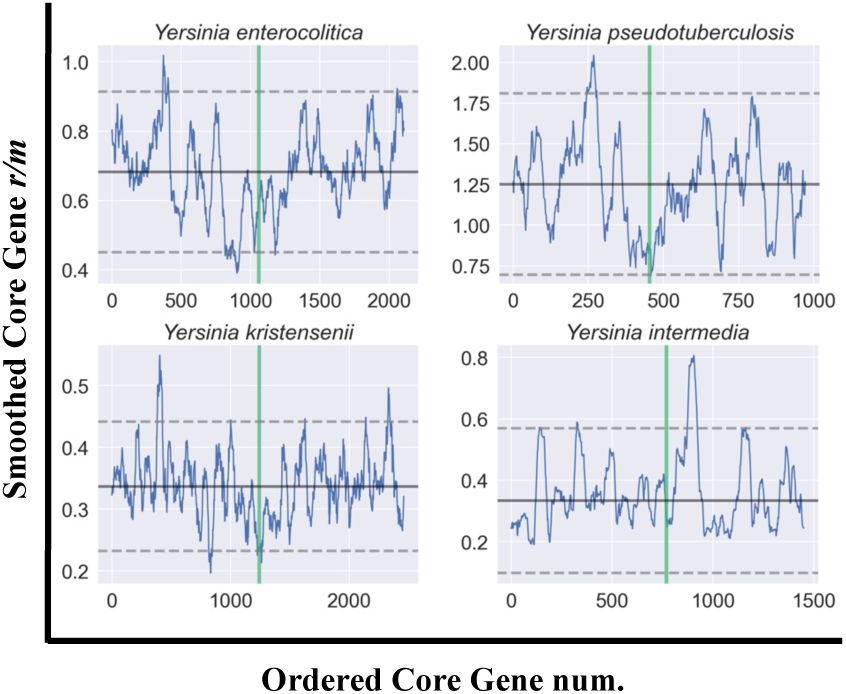
Landscape of recombination rate across *Yersinia* genomes (*n*=4). These graphs display the smoothed average across 50 estimable core genes with a step of 2 where *x* is the core gene number in order of its appearance with increasing distance from Ori. For each graph, *x*=0 demarcates Ori and the green line demarcates Ter. The black horizontal line denotes the average *r/m* value of all estimable core genes, and the dashed lines represent two standard-deviations from the mean *r/m*.

**Supplementary Figure 11.**
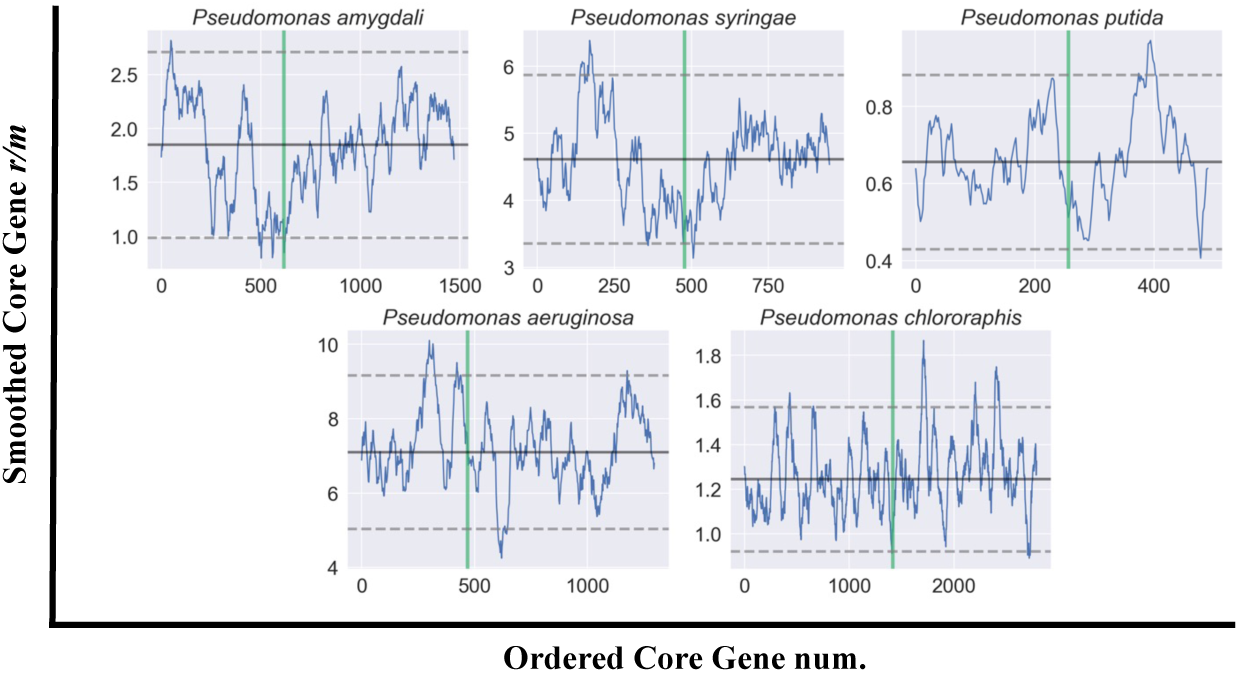
Landscape of recombination rate across *Pseudomonas* genomes (*n*=5). These graphs display the smoothed average across 50 estimable core genes with a step of 2 where *x* is the core gene number in order of its appearance with increasing distance from Ori. For each graph, *x*=0 demarcates Ori and the green line demarcates Ter. The black horizontal line denotes the average *r/m* value of all estimable core genes, and the dashed lines represent two standard-deviations from the mean *r/m*.

**Supplementary Figure 12.**
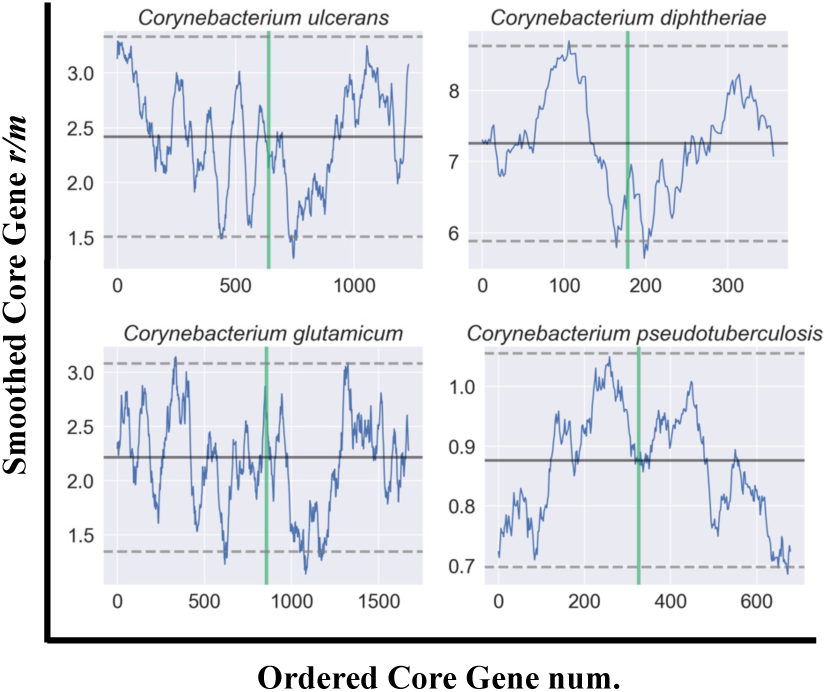
Landscape of recombination rate across *Corynebacterium* genomes (*n*=4). These graphs display the smoothed average across 50 estimable core genes with a step of 2 where *x* is the core gene number in order of its appearance with increasing distance from the Ori. For each graph, *x*=0 demarcates Ori and the green line demarcates Ter. The black horizontal line denotes the average *r/m* value of all estimable core genes, and the dashed lines represent two standard-deviations from the mean *r/m*.

## Supplementary Dataset Legends

Supplementary Dataset 1. Summary of gene data for each species analyzed in this study (*n*=146). The Dataset columns are as follows: Col 1) Species name, Col 2) Number of core genes (total), Col 3) Number of accessory genes, Col 4) Number of core genes without polymorphisms (*r/m* not estimable), Col 5) Number of core genes outside of simulation distribution (*r/m* not estimable), Col 6) Number of core genes used in this analysis (*r/m* is estimable), Col 7) Species’ core genome *r/m* (as per Torrance *et al*. (2024)), Col 8) Average *r/m* across estimable core genes, Col 9) Median *r/m* across estimable core genes, Col 10) Standard deviation of *r/m* across estimable core genes, Col 11) Average simulated *h/m* across estimable core genes, Col 12) Median simulated *h/m* across estimable core genes, Col 13) Standard deviation of simulated *h/m* across estimable core genes, Col 14) Average simulated *π* across estimable core genes, Col 15) Median simulated *π* across estimable core genes, Col 16) Standard deviation of simulated *π* across estimable core genes, Col 17) Average real *h/m* across estimable core genes, Col 18) Median real *h/m* across estimable core genes, Col 19) Standard deviation of real *h/m* across estimable core genes, Col 20) Average real *π* across estimable core genes, Col 21) Median real *π* across estimable core genes, and Col 22) Standard deviation of real *π* across estimable core genes.

Supplementary Dataset 2. Wilcoxon’s test comparison of *r/m* for core genes flanking accessory regions (“Flanking”) vs. *r/m* of core genes not flanking accessory regions (“Non-Flanking”) across all species *n*=146. Significant Benjamini-Hochberg adjusted *P*-values are highlighted in red. The median *r/m* of each group and difference in *r/m* between each group is also provided.

Supplementary Dataset 3. Spearman’s correlation test results for comparison between *r/m* and GC% across core genes for each species (*n*=146). Significant Benjamini-Hochberg adjusted *P*-values are highlighted in red.

Supplementary Dataset 4. Spearman’s correlation test results for comparison between *r/m* and *dN/dS* (Tab 1), *dN* (Tab 2), and *dS* (Tab 3) values across core genes for each species (*n*=142).

Supplementary Dataset 5. Spearman’s correlation test results for comparison between GC% and *dN/dS* values across core genes for each species (*n*=142). Significant Benjamini-Hochberg adjusted *P*-values are highlighted in red.

Supplementary Dataset 6. Wilcoxon test results for comparison of *r/m* between leading and lagging strand genes for each species (*n*=102). Significant Benjamini-Hochberg adjusted *P*-values are highlighted in red.

Supplementary Dataset 7. Spearman’s correlation test results for comparison of *r/m* values across the shared orthologs of related species (*n*=109 species pairs). Significant Benjamini-Hochberg adjusted *P*-values are highlighted in red. The Dataset also contains the pairwise divergence (branch length in A.A. substitutions per site) between each species pair.

Supplementary Dataset 8. Spearman’s correlation test results for comparison between *r/m* and distance from the Ori (Ori-Ter) and Ter (Ter-Ori) (*i.e.* both replichores) for *n*=102 species with circular chromosomes. Significant Benjamini-Hochberg adjusted *P*-values are highlighted in red.

## Availability of Data and Materials

All data used in this analysis was downloaded from NCBI’s Genbank public genomic repository. GenBank accession numbers and assembly IDs for all analyzed genomes are detailed in Supplementary Dataset 2 of Chapter 2. The pipeline *recABC* used to generate recombination rates is available at (https://github.com/lbobay/recABC). The datasets generated in this study (the core genomes, phylogenetic trees and the summary statistics of all the simulations for all species) are available on *Kaggle* at https://www.kaggle.com/datasets/ellistorr/bacterial-rm. The individual gene recombination rates, associated summary statistics, and other gene parameters used in this study are available at https://www.kaggle.com/datasets/ellistorr/Bacteria-Gene-rm.

## Funding Disclaimer

This report was prepared as an account of work sponsored by an agency of the United States Government. Neither the United States Government nor any agency thereof, nor any of their employees, makes any warranty, express or implied, or assumes any legal liability or responsibility for the accuracy, completeness, or usefulness of any information, apparatus, product, or process disclosed, or represents that its use would not infringe privately owned rights.

Reference herein to any specific commercial product, process, or service by trade name, trademark, manufacturer, or otherwise does not necessarily constitute or imply its endorsement, recommendation, or favoring by the United States Government or any agency thereof. The views and opinions of authors expressed herein do not necessarily state or reflect those of the United States Government or any agency thereof.

## Authors’ contributions

E.L.T., A.D., and L-M. B. designed, performed research, and analyzed data in this publication. E.L.T. and L-M.B. wrote this paper and A.D. reviewed and edited it.

## Acknowledgments

We would like to thank Ophelia Adams, Daniel Schrider and Kasie Raymann for providing advice and expertise. We would also like to thank Gavin Douglas, Hazem Sharaf, and Carolina Martinez-Gutierrez for their help in editing this manuscript.

